# Defining a critical enhancer near Nanog using chromatin-focused approaches identifies RNA Pol II recruitment as required for expression

**DOI:** 10.1101/2020.05.27.118612

**Authors:** Puja Agrawal, Steven Blinka, Kirthi Pulakanti, Michael H. Reimer, Sridhar Rao

**Affiliations:** Department of Cell Biology, Neurobiology, and Anatomy, Medical College of Wisconsin, Milwaukee, WI 53226 USA; Blood Research Institute, Versiti, Milwaukee, WI 53226 USA; Department of Pediatrics, Medical College of Wisconsin, Milwaukee, WI 53226 USA

**Keywords:** embryonic stem cells, super-enhancers, enhancers, gene expression

## Abstract

Transcriptional enhancers have been defined by their ability to operate independent of distance and orientation in plasmid-based reporter assays of gene expression. Currently, histone marks are used heavily to identify and define enhancers but both methods do not consider the endogenous role of an enhancer in the context of native chromatin. We employed a combination of genomic editing, single cell analyses, and sequencing approaches to investigate a *Nanog*-associated *cis*-regulatory element (CRE) which has been reported by others to be either an alternative promoter or a super-enhancer (SE). We first demonstrate both distance and orientation independence in native chromatin, eliminating the issues raised with plasmid-based approaches. We also demonstrate that the dominant SE modulates *Nanog* globally and operates by recruiting and/or initiating RNA Polymerase II. Our studies have important implications to how transcriptional enhancers are defined and how they regulate gene expression.

**AUTHOR SUMMARY:** Different DNA elements help regulate the levels of gene expression. One such element are enhancers, short sequences that interact with genes to modulate levels of expression but can operate over large distances. Previously, these sequences were defined by their ability to regulate expression independent of their distance from a gene and the orientation of the sequence. However, these characteristics were found using techniques that did not recapitulate the native environment. Here, we have shown that an enhancer of one gene is indeed an enhancer by testing its distance and orientation-independence within the native environment. We also show that the mechanisms by which the enhancer is regulating expression is by controlling the levels of RNA Polymerase II at a gene. RNA Polymerase II is the protein that converts the gene sequence to a form usable by a cell, called mRNA. This is interesting because while this has been considered historically the main way enhancers operate, more recent work has focused on other, later regulatory steps involved in controlling mRNA production.

## INTRODUCTION

Gene expression is regulated by two types of genetic elements: *Trans* elements typically encode proteins such as transcription factors (TFs) which subsequently bind *cis* regulatory elements (CREs) that must be on the same DNA molecule as the gene they regulate. Different types of CREs have historically been classified based upon their behavior in plasmid-based reporter assays [1,2]. For almost 40 years it has been accepted that promoters are required to be in the correct orientation and immediately adjacent to the gene they regulate whereas enhancers operate independent of both distance and orientation. The advent of enhancer-specific epigenetic signatures based on histone marks such as H3K27Ac or H3K4me1 permit genome-wide identification of enhancers which then demonstrate enhancer activity in reporter assays [3,4]. However, plasmid-based assays are limited for multiple reasons. First, they do not fully recapitulate native chromatin. Second, they typically are performed on smaller (<500bp) DNA sequences rather than the larger chromatin domains of many highly active enhancers. Third, they cannot precisely link a given enhancer sequence to the gene(s) they may regulate *in vivo*. As such, plasmid assays are far more effective at confirming DNA sequences with enhancer potential, rather than definitively identifying them as enhancers.

The advent of sequencing based chromosomal conformation capture techniques has allowed the measurement of genome-wide enhancer-gene interactions [5,6]. Interestingly, this approach demonstrates that many enhancers interact with multiple genes and vice versa, but are insufficient to properly determine if an enhancer is required for gene(s) expression [7,8]. The classic approach to address this question is through genetics, namely deleting a putative enhancer and measuring mRNA levels of nearby genes, a method made highly feasible through genomic editing approaches such as CRISPR/Cas9. One important point is that while these approaches can identify which gene(s) are regulated by an enhancer, many of the mechanistic details of how the enhancer regulates transcription to modulate gene expression are not elucidated through solely this approach.

Multiple models of enhancer-mediated gene expression exist within the literature. Early theories postulated that enhancers looped in to interact with promoters and recruited RNA Polymerase II (RNAPII) to the gene’s promoter (Rippe et al., 1995). More recently, multiple mechanisms have been proposed which focus on enhancers regulating transcriptional elongation, including promoter-proximal pause release of RNAPII through various mechanisms (reviewed in Chen et al. 2018). It has also been proposed that enhancers modulate transcriptional bursting, or the periods of time during which transcription is active, which represents a combination of initiations and elongation [11,12]. New studies demonstrate that among enhancers there is a sub-class of highly active enhancers called “super-enhancers” (SEs; Pulakanti et al., 2013; Whyte et al., 2013), which can form phase separated droplets within the nucleus to concentrate transcriptional machinery around highly transcribed genes [15]..

The extended *Nanog* locus is a unique locus to study how super-enhancers regulate gene expression and pluripotency. The *Nanog* locus (150kb) contains a number of different pluripotency-associated genes including *Dppa3, Gdf3*, and *Apobec1* [7,16]. It also contains three SEs (−5, −45, and +60, based upon distance in kb from TSS) which interact with *Nanog* and behave as enhancers in reporter assays [7]. One group has argued that the −5 SE is actually an alternative promoter, emphasizing that plasmid-based approaches are insufficient to determine if a DNA element is a promoter or enhancer [17]. In this study we demonstrate the −5 SE is an enhancer by confirming it operates in a distance and orientation independent fashion through genomic approaches and regulates *Nanog* by modulating RNAPII initiation or recruitment.

## RESULTS

### The −5 *Nanog* CRE is required for ESC pluripotency in a Nanog-dependent manner

In our prior study [7] we were unable to recover mouse embryonic stem cells (ESCs) which exhibited biallelic deletion of the −5 *cis* regulatory element (CRE), leading us to hypothesize it is required for pluripotency. We refer to this element as a CRE rather than a super-enhancer (SE) because one group has reported this element is an alternative promoter [17]. To identify if this element is required for *Nanog* expression, we used genomic editing to insert a tamoxifen (4OHT) inducible Cre-recombinase (CreER^T2^) into the constitutively expressed *Rosa26* locus in ESCs to facilitate conditional deletions, and then biallelically inserted *loxP* sites to flank a 2.5 kb region of the −5 CRE to encompass two Nanog, Oct4, and Sox2 (NOS) binding sites (Supplemental Fig 1A, Fig 1A, left). Treatment with 4OHT induces complete biallelic deletion of the −5 CRE as compared to vehicle treated (ethanol; Supplemental Fig 1B, Fig 1A, left). ESCs began to differentiate and became non-adherent, consistent with a loss of pluripotency, following 4OHT exposure. Staining for the pluripotency marker alkaline phosphatase was reduced in cells treated with 4OHT compared to control (Supplemental Fig 1C-i,ii). Deletion of the −5 CRE resulted in a rapid loss of Nanog mRNA (Fig 1A, right) and protein (Supplemental Fig 1D). By contrast, *Gdf3*, a nearby gene, showed virtually no change in expression following deletion of the −5 CRE (Fig 1B). There was also a decrease in other pluripotency-associated TFs such as *Oct4, Esrrb*, and *Klf4*, demonstrating a progressive collapse of the transcriptional network regulating pluripotency (Fig 1B). Consistent with previous studies showing that Nanog represses endoderm specification, RT-qPCR for key differentiation genes (Fig 1C) and endoderm-promoting TFs such as *Gata4, Gata6* and *Hnf4a* (Fig 1D) demonstrated increased expression following 4OHT treatment. These results demonstrate that the −5 CRE is required for ESC pluripotency, likely by regulating *Nanog* expression.

**Fig 1.**
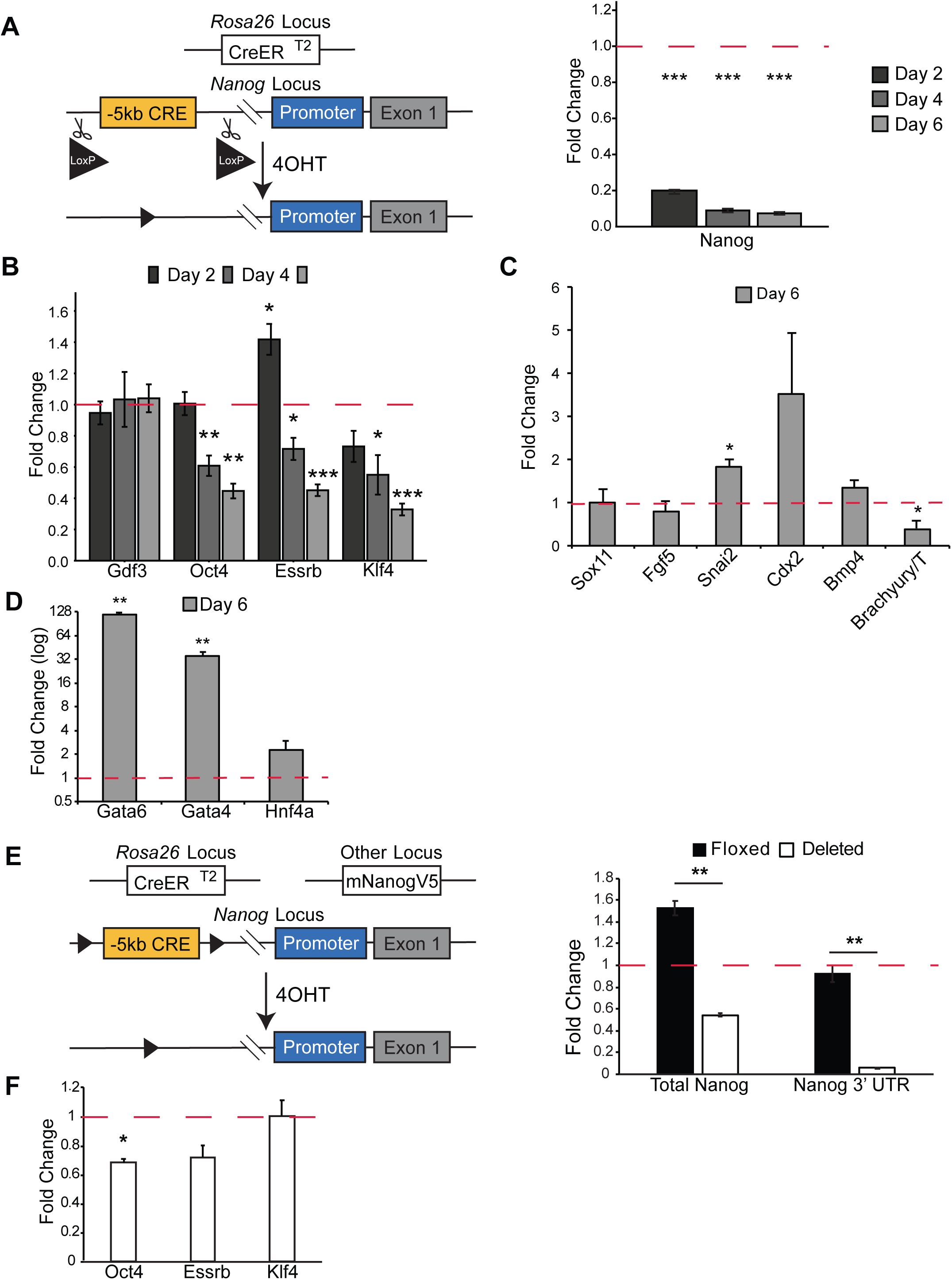
Deletion of the −5 CRE. (A) Biallelic deletion of −5 CRE was achieved by inserting two loxP sites around the enhancer and cells were treated with tamoxifen (4OHT) for six days. Left panel, schematic; right panel, mRNA levels. (B) Expression for *Gdf3* and pluripotency markers for cells in (A) treated with 4OHT for 6 days (C) Day 6 mRNA expression of mesodermal, ectoderm, and trophectoderm differentiation markers. (D) Day 6 mRNA expression of endodermal promoting transcription factors. (E) Stable biallelic deletion of the −5 CRE was achieved by rescuing with a mouse *Nanog* cDNA. Endogenous *Nanog* expression is measured via the *Nanog* 3’UTR, while Total Nanog measured endogenous and the exogenous expression. Left panel, schematic; right panel, mRNA levels. (F) Pluripotency markers in stably −5 CRE deleted cells shown in Fig 1B. All mRNA levels measured by RT-qPCR, shown relative to wildtype (dotted red line) *p<0.05, **p<0.01, ***p<0.001 Student’s One Sample t-test.

Next, we hypothesized that the −5 CRE maintains pluripotency solely by regulating *Nanog* expression rather than the expression of another gene on chromosome 6 (chr6). To test this, we made a stable cell line expressing murine *Nanog* with a ubiquitous promoter (CAG; Fig 1E, left). Prior to 4OHT treatment *Nanog* mRNA levels are approximately 50% higher than wild-type ESCs, which then falls following 4OHT treatment to 50% below wild-type (Fig 1E, right). Importantly, *Nanog*^*+/-*^ animals are viable and ESCs remain pluripotent [18,19], indicating that 50% levels of Nanog do not compromise pluripotency. Endogenous *Nanog* gene expression can be followed with RT-qPCR primers amplifying the *Nanog* 3’UTR which is absent from the *Nanog* transgene. Post-4OHT treatment these cells show a profound (>90%) reduction in endogenous *Nanog* expression (Fig 1E, right) and a small decrease in *Oct4* levels but no significant change in other core pluripotency TFs such as *Esrrb* or *Klf4* (Fig 1F), and a rescue of the alkaline phosphatase phenotype (Supplemental Fig 1C-iii, iv) demonstrating the loss of pluripotency in cells without the *Nanog* transgene is attributable to the loss of *Nanog* expression.

To determine if the −5 CRE is solely regulating *Nano*g we used RNA-seq to identify other altered transcripts. First, we identified genes on chr6 which showed at least a 2-fold, statistically significant change (adj p-value<0.05) between samples (Fig 2A) and as a control compared to data where *Nanog* was depleted by RNAi [20]. For comparison we queried changes on chr5&7 to estimate gene expression changes secondary to *trans* effects from changes in Nanog protein levels. None of the altered genes were within 1MB of *Nanog* except *Dppa3*, which was increased in expression as expected since it is directly repressed by Nanog protein binding to its promoter [7]. To further clarify if loss of the −5 CRE affected the expression of other genes *in cis* we queried expression changes of all genes within the *Nanog* Topologically Associated Domain (TAD) as well as the two adjacent TADs [21], irrespective of statistical significance or fold-change (Fig 2B). With the exception of *Nanog* and *Dppa3*, most genes showed minimal gene expression changes and were similar to those when *Nanog* was depleted by RNAi, implying this was due to reduced Nanog protein operating in *trans*. We therefore conclude that the −5 CRE exclusively regulates *Nanog* expression, with no evidence it regulates other genes on chr6 in *cis*. Collectively, these experiments demonstrate that the *Nanog* −5 CRE is required for pluripotency through its direct regulation of *Nanog* expression.

**Fig 2.**
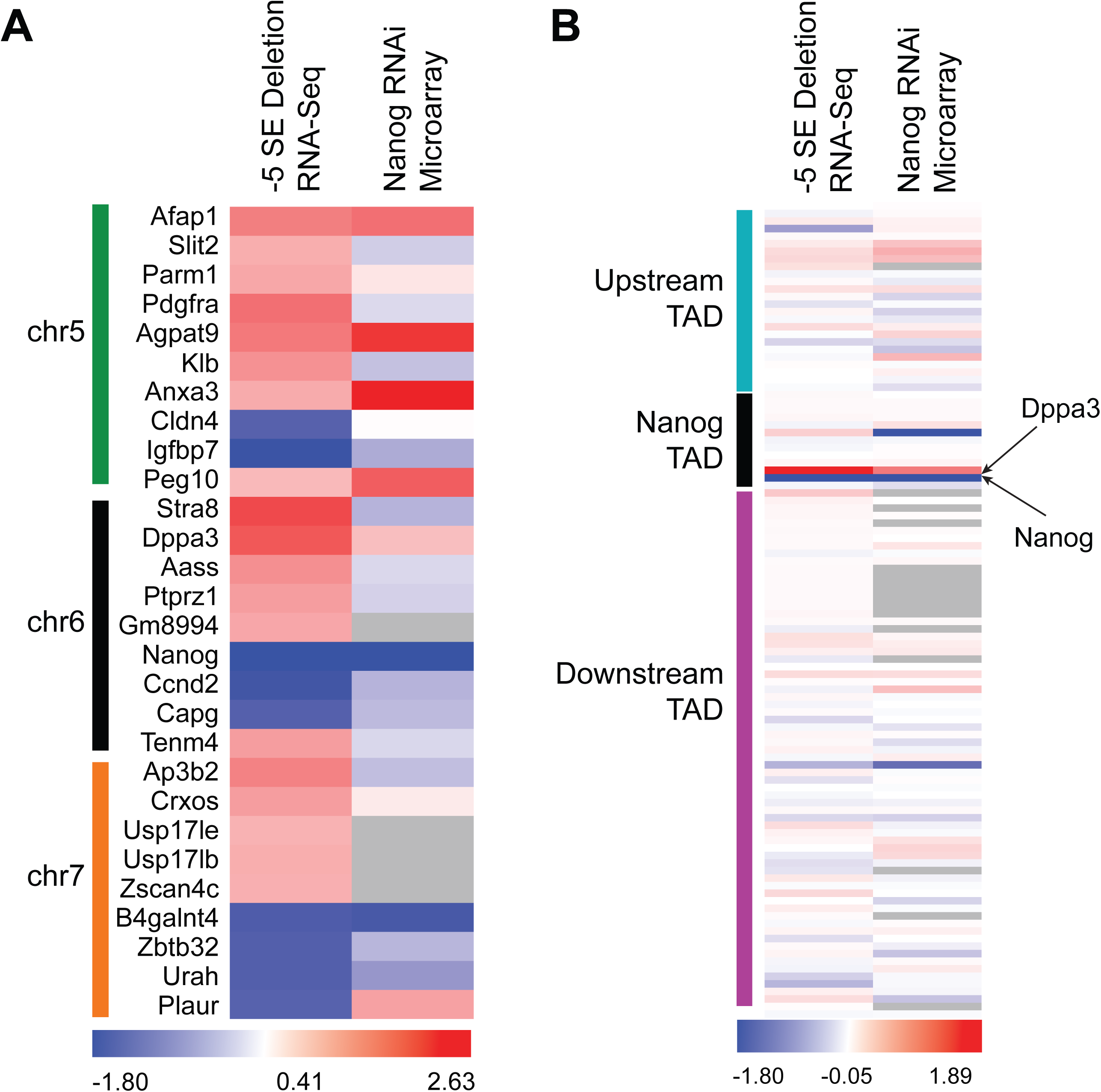
The −5 CRE regulates *Nanog* exclusively. (A) Differentially expressed genes, defined to be at least two-fold, statistically significant (adj p<0.05) change, on chromosome 5,6, and 7 via RNA-seq. *Nanog* RNAi microarray data is shown for comparison^20^. (B) Differentially expressed genes, irrespective of significance or fold-change, on chromosome 6 within the *Nanog* TAD, one TAD upstream and one downstream.

### The *Nanog* −5 CRE operates in a distance and orientation independent fashion

The *Nanog* −5 *cis*-regulatory element (CRE) in plasmid assays acts independent of distance and orientation and has been extensively referred to as an enhancer within the literature [22,23]. By contrast, in at least one report the −5 CRE was considered an alternative promoter which played a critical role in regulating pluripotency through an alternative *Nanog* isoform [17]. Given this ambiguity we chose to definitively establish if this element had enhancer activity within the context of normal chromatin with our 4OHT-inducible Cre-LoxP system by inserting one of the *LoxP* sites in the opposite orientation (Fig 3A, left). In this configuration, Cre activation by 4OHT treatment will induce biallelic “flipping” of the −5 CRE continuously between the two orientations. To isolate clones with biallelic inversion of the −5 CRE we grew the cells at clonal density and isolated individual colonies following 4OHT treatment and verified inversion of the −5 CRE by PCR (Supplemental Fig 2A). Comparing the biallelic inversion to wild-type orientation of the −5 CRE we observed no statistically significant changes in the expression of *Nanog, Dppa3, Oct4, Esrrb*, or *Klf4* (Fig 3A, right). To verify there was no change in the Nanog protein banding pattern, we performed western blots and did not observe loss of a band which may represent an alternative isoform (Supplemental Fig 2B). This data is consistent with the −5 CRE regulating *Nanog* expression in an orientation independent manner, a classic property of an enhancer but not a promoter. While we cannot rule-out that this CRE is simultaneously acting as both an enhancer and an alternative promoter, our data demonstrates that any alternative isoforms produced via the −5 CRE as a promoter [17] are dispensable for pluripotency.

**Fig 3.**
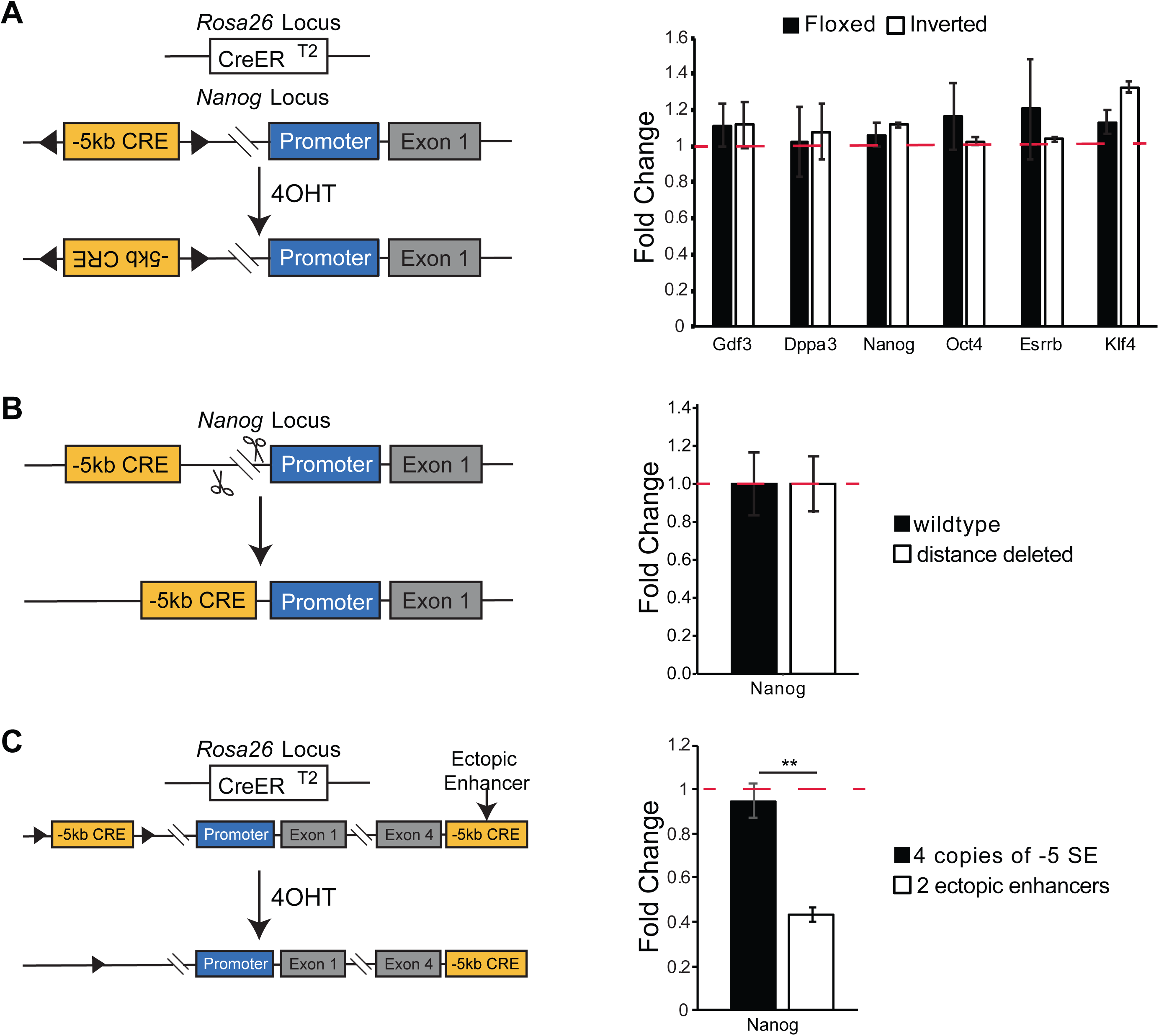
Manipulation of the −5 CRE. (A) The −5 CRE was flipped by inserting opposing loxP sites and then treatment with tamoxifen. Left panel, schematic; right panel, mRNA levels of *Nanog* and relevant pluripotency genes. (B) An ≈2 kb region between the −5 CRE and *Nanog* TSS was deleted. Left panel, schematic; right panel, *Nanog* mRNA levels. (C) Two copies of the −5 CRE were inserted downstream of *Nanog* in cells where the endogenous enhancer is floxed. Endogenous enhancers were deleted via treatment with tamoxifen. Left panel, schematic; right panel, mRNA levels of *Nanog*. All mRNA levels measured by RT-qPCR, shown relative to wildtype (dotted red line) *p<0.05, **p<0.01, ***p<0.001 Student’s One Sample t-test.

We next hypothesized that the −5 CRE would also operate independent of distance from the *Nanog* transcriptional start site (TSS). To determine this, we first deleted the intervening ≈2 kb between the - 5 CRE and promoter (Supplemental Fig 2C, Fig 3B, left) and found no significant change in *Nanog* expression (Fig 3B, right). Next, we biallelically inserted an additional copy of the −5 CRE between the *Nanog* TES and the nearest CTCF site to ensure it was in the same insulated neighborhood (Supplemental Fig 2D-E, Fig 3C, left; Dowen et al., 2014). Insertion of the additional −5 CRE caused no change in *Nanog* mRNA levels (Fig 3C, right). Treatment with 4OHT resulted in deletion of the endogenous −5 CRE and caused a reduction in *Nanog* mRNA by approximately 50% (Fig 3C). Expression of other key pluripotency markers such as *Oct4, Esrrb*, and *Klf4* were unchanged, indicating that pluripotency was maintained (data not shown). The rescue of *Nanog* expression by the ectopic enhancers is consistent with the −5 CRE operating independent of distance, albeit less effectively then its native chromatin position. One reasonable explanation for the reduced *Nanog* expression with the ectopic enhancer is that the −5 SE includes a larger chromatin domain, whereas we inserted only the core ≈2.5kb containing two NOS sites into an alternative location. This may imply that additional sequences surrounding the core are required for full activation. Alternatively, moving the enhancer further away may prevent it from fully activating Nanog. Nonetheless, this demonstration of enhancer function represents a highly feasible, native chromatin approach to confirm a DNA element is an enhancer. Collectively, these data demonstrate that the −5 CRE is an enhancer and will hereafter be referred to as the −5 SE [14].

### Individual sub-enhancers within the −5 super-enhancer are additive in regulating *Nanog* expression

Recent work from our group and others has demonstrated that the −5 SE is a super-enhancer based upon several criteria, including high levels of the epigenetic mark H3K27Ac, robust binding by the Mediator complex, and production of an enhancer-transcribed RNA (eRNA; Pulakanti et al., 2013; Whyte et al., 2013). Several groups have demonstrated that within a single super-enhancer a single smaller sub-enhancer is required for its function. By contrast, the remaining sub-enhancers are considered dispensable for regulating gene expression [25]. To determine if the −5 SE had a dominant sub-enhancer, we first reviewed published chromatin immunoprecipitation coupled with next generation sequencing (ChIP-seq) datasets from other groups to determine if there were sub-enhancers within the larger −5 SE. We observed there were two distinct regions occupied by the classic pluripotency transcription factors Nanog, Oct4, and Sox2 (Fig 4A). To determine if one, or both sub-enhancers were critical to pluripotency we deleted each individually with CRISPR/Cas9 using a pair of distinct gRNAs (Fig 4A-B, Supplemental Fig 2F). Surprisingly, we were able to recover biallelic deletion of the individual 5’ or 3’ sub-enhancers without difficulty. Deletion of either the 5’ or 3’ sub-enhancer results in an approximately 50% reduction in *Nanog* mRNA, but these reductions were not sufficient to alter pluripotency as measured by *Oct4, Esrrb, or Klf4* expression (Fig 4C). This demonstrates that neither the 5’ nor the 3’ sub-enhancer is required for pluripotency even though both are required as part of the −5 SE for normal *Nanog* expression. This is likely consistent with locus-specific issues surrounding how enhancers regulate gene expression, making it likely difficult to generalize findings across all enhancers. Alternatively, it may be the two sub-enhancers function in an additive fashion. Nonetheless, it confirms that these two sub-enhancers at the heart of the −5 SE are collectively required for proper *Nanog* expression.

**Fig 4.**
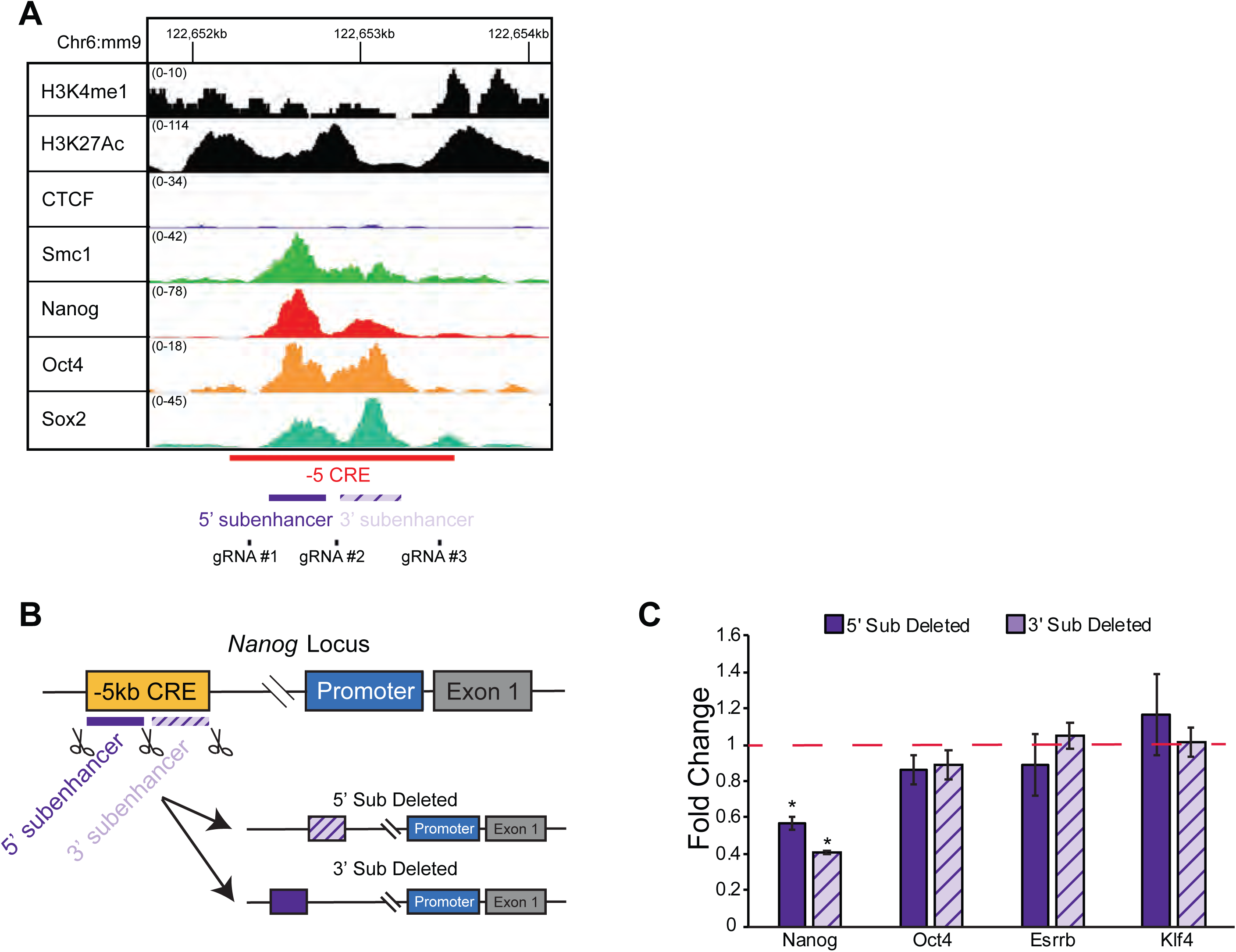
-5 SE sub-enhancers are both required. (A) IGV snapshot of the −5 CRE showing two sub-enhancers and locations of guide RNAs. X-axis is genomic position, Y-axis is normalized tag count. (B) Schematic (C) mRNA levels of *Nanog* and relevant pluripotency genes when two sub-enhancers within the −5 CRE were deleted. RT-qPCR data is shown relative to wildtype (dotted red line) *p<0.05, **p<0.01, ***p<0.001, Student’s One Sample t-test.

### -5 SE regulates expression in all cells

Previous studies from our lab demonstrated that monoallelic deletion of the −5 SE, biallelic deletion of the −45 SE and biallelic deletion of the +60 SE have different effects on *Nanog* expression despite each enhancer physically interacting with the gene, as shown by Chromosomal Conformation Capture (3C; Blinka et al., 2016). Specifically, deletion of the −45 SE causes a 50% decrease in *Nanog* expression, while deletion of the +60 SE had no change in *Nanog* expression. Work from this study has further shown that the −5 SE is critical to *Nanog* expression, as there is a 90% decrease in *Nanog* expression upon biallelic deletion (Fig 1E, right). This has led us to question how two enhancers could both be required for maximal *Nanog* expression, since most recent literature argues that at any given time point a single enhancer operates on a gene [11]. We hypothesized that each enhancer may operate on distinct sub-populations of cells, which we could not distinguish using a bulk population of cells. Specifically, the −5 SE could be regulating a larger proportion of high *Nanog* expressing cells, while the −45 SE and +60 SE regulate separate, low-expressing cells (Supplemental Fig 3A-C).

To investigate this possibility, we performed single-cell RT-qPCR on the −5 SE biallelically deleted cells with *Nanog* in *trans* (Fig 1E). If the −5 SE is regulating a different population of cells, we should observe a bimodal distribution of *Nanog* expression following enhancer deletion (Supplemental Fig 3B). By contrast, if the −5SE is operating on *Nanog* in all cells, we should observe a uniform reduction in *Nanog* mRNA levels (Supplemental Fig 3C). It should be noted that although some groups have shown that *Nanog* has bimodal expression in single cells [26], the presence of truly bimodal *Nanog* expression remains an open debate [27,28]. Interestingly, we found a uniform reduction in endogenous *Nanog* expression as measured by the 3’UTR when deleting either the −5 or −45 SE (Fig 5). Two controls, *Oct4* and *ERCC3*, showed minimal changes in expression. Collectively, these single cell experiments support a model that the −5 SE is the dominant enhancer in all cells, and even with its deletion neither the −45 nor +60 SEs can compensate.

**Fig 5.**
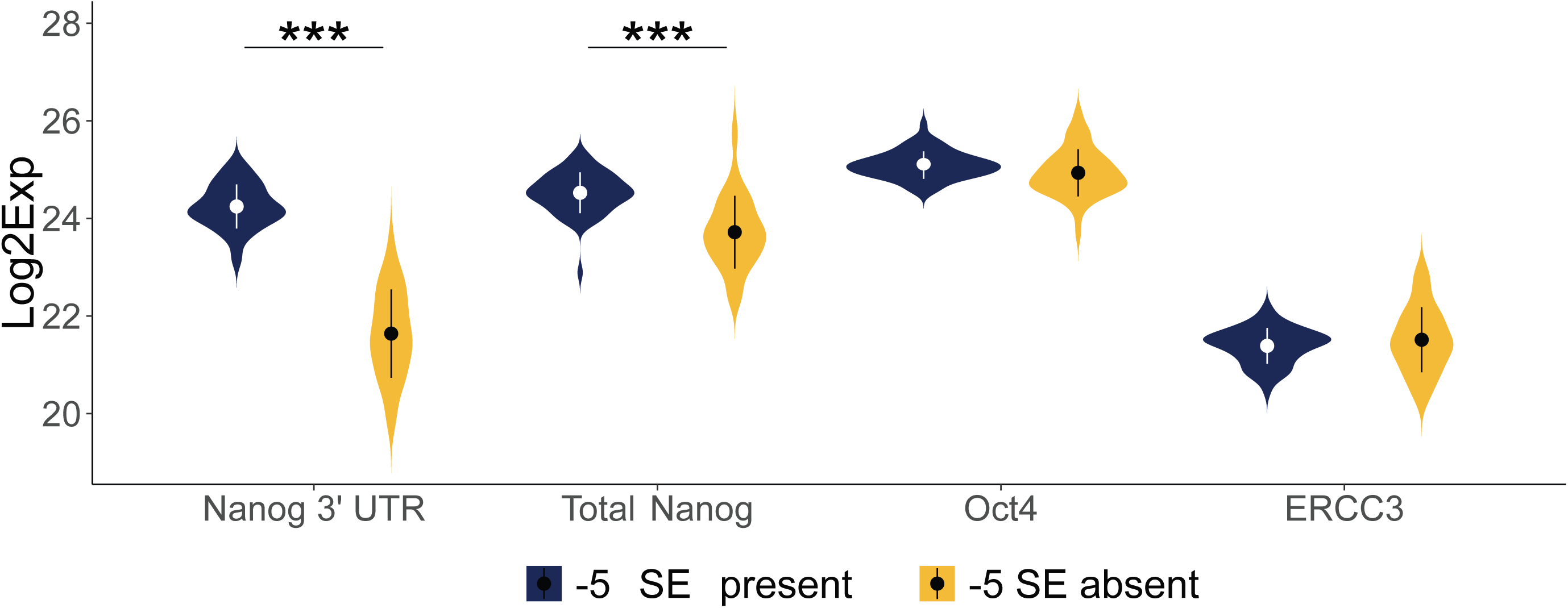
-5 SE operates in all cells. Single-cell RT-qPCR for endogenous *Nanog* via *Nanog* 3’UTR, Total *Nanog, Oct4*, and *ERCC3* in −5 SE deleted cells. Expression is depicted as Log2Exp relative to the Limit of Detection, as described by Fluidigm. ***p<0.001 by Mann-Whitney test.

### The −5 SE regulates *Nanog* by regulating transcriptional initiation/recruitment

Previous studies have argued that SEs regulate gene expression through promoter-proximal pause release (hereafter referred to as pause release) of RNAPII [29,30]. Briefly, transcription begins with the recruitment of RNAPII to the promoter, which is immediately phosphorylated on Ser 5 (Ser5P) of its c-terminal domain (CTD), which results in bidirectional transcription around the TSS and is referred to as “paused” RNAPII because it cannot elongate further into the gene body. The “pausing” of RNAPII occurs 20-120bp downstream of the TSS, which must be relieved for productive gene transcription [10]. “Pause release” is mediated by phosphorylation of Ser2, releasing RNAPII to transcribe the gene body and is referred to as elongating RNAPII. To measure changes in RNAPII dynamics, we performed CUT&Tag [31] with antibodies specific to total and paused (RNAPII-Ser5P) RNAPII in WT and −5 SE deleted cells.

Depending on which phase of transcription an enhancer is regulating, RNAPII’s genomic location will change as shown in Supplemental Fig 4. As described above, RNAPII is phosphorylated on the Ser5 position of its CTD after recruitment, at which point it is paused. If recruitment is regulated by the enhancer, loss of the enhancer will cause a loss RNAPII-Ser5P at the TSS (Supplemental Fig 4-i). If pause release is being regulated, loss of an enhancer will cause a build-up of RNAPII-Ser5P that cannot be released (Supplemental Fig 4-ii). If neither of these are the steps being regulated by the enhancer, RNAPII-Ser5P enrichment will remain unchanged (Supplemental Fig 4-iii). An inherent flaw of this system is that RNAPII binding at *Nanog* exons in the 0-copy cell line is confounded by the presence of the *Nanog* cDNA in *trans*. We observed a complete loss of paused RNAPII in the 0-copy cell line (Fig 6A), consistent with the −5 SE playing a critical role in RNAPII recruitment and/or phosphorylation on Ser5. These data indicate that the −5 SE regulates *Nanog* not through RNAPII pause-release, but rather through modulating transcriptional initiation/recruitment.

**Fig 6.**
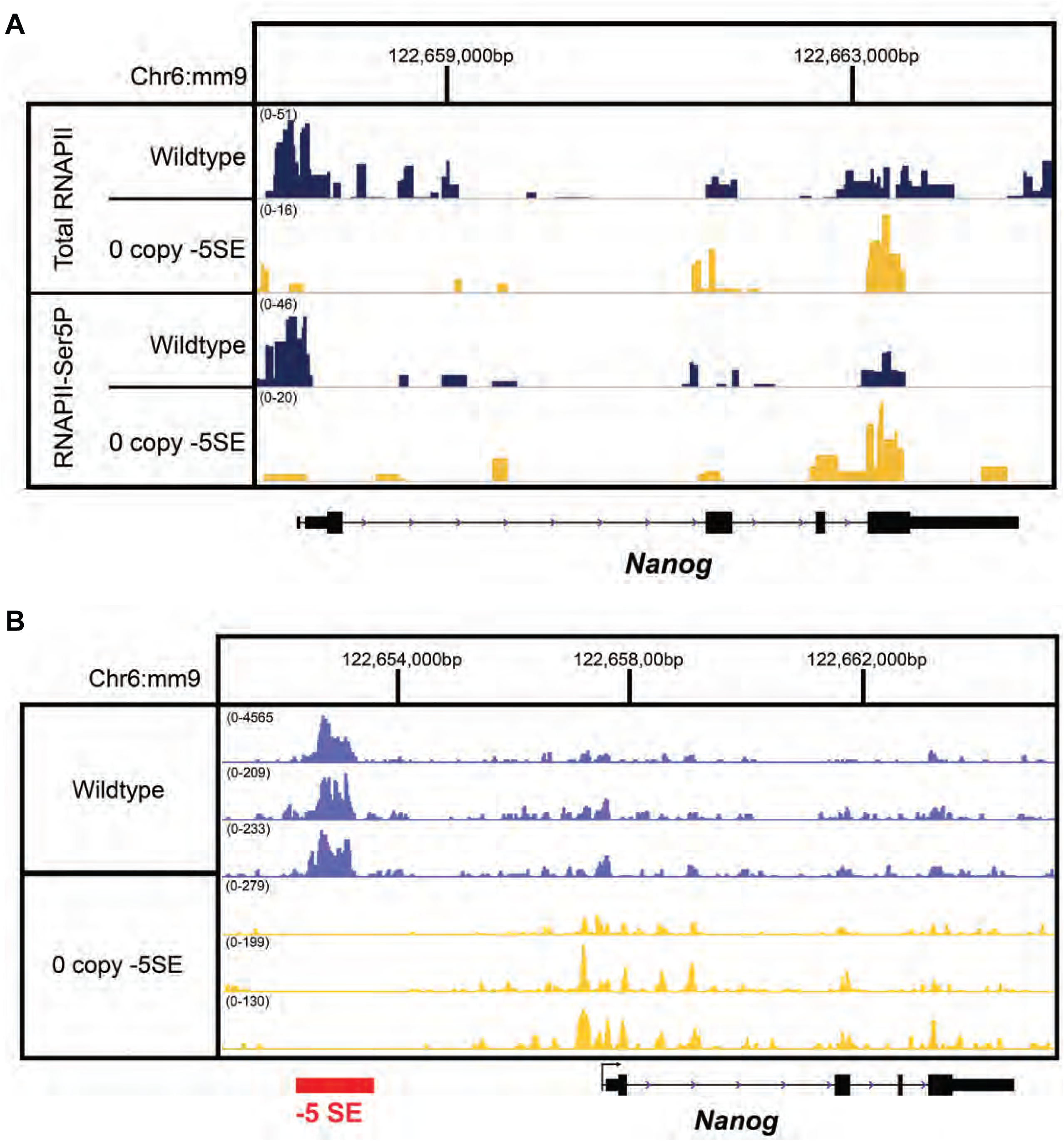
-5 SE operates prior to RNAPII pause release. (A) IGV snapshot of CUT&Tag for Total RNAPII (n=1) and RNAPII-Ser5P (n=3) in wildtype and 0 copy of −5 SE cells. Note that exonic regions of the 0 copy cell line are confounded by the exogenous *Nanog* cDNA. (B) IGV snapshot of ATAC-Seq data, separated by each sample of each wildtype cells where the −5 SE is floxed and cells with the −5 SE completely deleted. Genes and SEs are shown below. X-axis is genomic position; Y-axis is normalized read count.

These data led us to question whether the −5 SE is modulating the initial binding of RNAPII to *Nanog* by changing chromatin accessibility, rather than directly recruiting RNAPII. To investigate this further, we performed assay for transposase-accessible chromatin using sequencing (ATAC-seq) on wildtype cells and −5 SE deleted cells (Fig 1E) and found no significant differences in chromatin accessibility within the *Nanog* locus (Fig 6B). Thus, we conclude that the mechanism by which the −5 SE regulates *Nanog* is by modulating RNAPII recruitment/initiation but not through a change in chromatin accessibility. Although we cannot distinguish between transcriptional initiation versus recruitment, given the complete loss of RNAPII at *Nanog* upon deletion of the −5 SE, recruitment of RNAPII to the promoter is likely the rate-limiting step modulated by the enhancer, since generation of the initiating form of RNAPII (Ser5p) is not rate limiting. Together, these data show that the −5 SE is an enhancer critical to *Nanog* expression prior to pause-release of RNAPII.

## DISCUSSION

While enhancers have been well-known regulators of gene expression for 40 years, it has become apparent with new technologies that they are far more numerous then classical protein coding genes (>4 fold) and unlike promoters far more variable across tissues, implying enhancers play a central role in regulating tissue-specific expression [32–34]. Given their importance, the breadth of questions which remain within the field is profound. In particular, the reliance on plasmid-based approaches have been *de rigeur* for the formal definition of whether a *CRE* has enhancer activity. For the −5 SE, given its relative proximity to the *Nanog* TSS and literature suggesting it may have a promoter like activity [17], plasmid-based assays alone were unlikely to definitively address that issue. Importantly, this DNA element may function as an alternative promoter in other context, such as alternative pluripotent states or primordial germ cells, both of which express *Nanog* [35]. Given the ease of genetically engineering ESCs using CRISPR/Cas9 technology, our initial goal was to move beyond plasmid-based approaches and utilize native cells/chromatin to determine whether the −5 SE truly displayed enhancer potential. No different than a plasmid, we utilized a combination of classical genetic approaches. First, the −5 SE is required for pluripotency which can be genetically rescued by supplying Nanog *in trans*. Second, reversing the orientation of the −5 SE had no effect on pluripotency nor *Nanog* expression. Third, moving the enhancer either closer to the *Nanog* TSS or within the insulated neighborhood but downstream of the TSS permitted sufficient *Nanog* expression to maintain pluripotency. Importantly, this conclusively demonstrates the −5 SE is an enhancer, but with two caveats. First, while the −5 SE is an enhancer, these studies does not rule-out that it is also a promoter, as suggested by others [17]. Reversing the orientation of the −5 SE had no effect on *Nanog* expression and pluripotency, this formally demonstrates that if it does operate as a promoter, this aspect of the −5 SE function is not formally required for pluripotency. Second is that insertion of the −5 SE downstream of the *Nanog* TES did not completely recapitulate normal expression. One reasonable explanation is that the −5 SE includes a larger chromatin domain, whereas we inserted the core ≈2.5kb into an alternative location. This may imply that additional sequences surrounding the core are required for full activation when it is moved into a new location. Alternatively, it may be that simply moving the enhancer father away effects its ability to fully activate Nanog expression. Nonetheless, this demonstration of enhancer function represents a highly feasible, native chromatin approach to assess enhancer function.

By the end of our studies we realized the *Nanog* locus has essentially been converted to a reporter gene, with minimal alteration beyond those described and the insertion of *LoxP* sites. This minimizes confounding variables based upon artificial reporters such as the extensive presence of bacterial DNA sequences and/or insertion into a heterologous region of the genome. In addition, because *Nanog* is critical to ESCs, it permitted additional experiments to understand how the enhancer may modulate pluripotency and/or differentiation. One caveat is that many of our experiments were performed by supplying Nanog *in trans*, thereby permitting the cells to maintain pluripotency even when native *Nanog* expression was significantly reduced by deleting the −5 SE. Given our use of a heterologous, constitutively active promoter (CAG) and a *Nanog* cDNA which lacked the 5’ UTR, we can’t ensure that subtle changes in the temporal regulation of *Nanog* expression or Nanog protein levels were preserved. Nonetheless, supplying Nanog *in trans* was able to restore approximately heterozygous levels of *Nanog*, which is sufficient to maintain pluripotency and substantially suppress spontaneous differentiation of ESCs [18,19].

Recent literature has shown there are many mechanisms by which enhancers regulate gene expression. It was previously thought that enhancers promoted the recruitment of RNAPII and other transcriptional machinery to the promoter to “promote” transcription [9]. Recent studies focusing on highly active enhancers have focused on their role in RNAPII pause release, the first steps of RNAPII converting from the initiating to the elongating form [29,36]. Multiple studies have also shown that enhancers regulate transcriptional bursting [11,12], the observation of oscillating transcriptional activity over time, in which one transcriptional burst is a period of time during which there is active transcription. Enhancers have been shown to regulate the frequency of these bursts of activity, which represent a combination of initiation and elongation. More recently, enhancers have been implicated in forming phase-separated condensates that concentrate transcriptional machinery for actively transcribed genes [15,37,38]. Critically, these studies have not demonstrated how multiple enhancers could simultaneously regulate expression of a single gene, and if enhancers uniformly operate through the same or different mechanisms on the same gene. Since the −5 SE is indispensable for *Nanog* expression, and cannot be compensated for by another enhancer, the obvious question is whether this is because the enhancers function through different mechanisms, or perhaps in other pluripotent states. Deletion of each enhancer at the locus causes variable changes in *Nanog* expression [7], further leading us to ask if each enhancer plays a unique role in regulating *Nanog* expression through different phases of transcription. For example, it may be that while the −5 SE does not regulate *Nanog* through pause-release, another enhancer plays that more traditional role when modulating *Nanog* expression.

Given the broad role of enhancers in regulating tissue-specific gene expression, our work has implications for how other gene:enhancer pairs are studied. In the absence of genetic confirmation, it is difficult to confirm an enhancer:gene functional dyad based solely on plasmid-based approaches. In addition, further attention needs to be paid to the other enhancers in the region to understand how multiple enhancers work together to regulate a gene. Understanding the interplay of the three SE’s around *Nanog* will further drive changes in how gene:enhancer pairs are studied, especially since they may operate through different phases of transcription to regulate expression.

## METHODS

For further information and requests for reagent and resources, please contact the Lead Contact, Sridhar Rao (; 414-937-3841).

### Cell culture

Gelatin-adapted ESCs were utilized for all experiments. These are male, in-house generated, ICM-derived 129SVJ derived murine ESC line, similar to the one we have used previously and cultured under similar Serum/LIF conditions[39] [40]. Briefly, cells were propagated under feeder-free conditions in DMEM (Corning #10-017-CV) with the following supplements (FBS – GemBio #100-106, Penicillin/Streptomycin – Corning #30-002-Cl, MEM Nonessential Amino Acids – Corning #25-025-Cl, L-glutamine – Corning #25-005-Cl, Nucleosides – Sigma #ES-008-D, LIF, β-mercaptoethanol at the appropriate concentration). 2 µM of 4-Hydroxytamoxifen (4OHT) in 70% ethanol (EtOH) was used for all experiments and diluted approximately 1:1000 for drug treatments with EtOH as a control.

### CRISPR/Cas9 mediated genomic editing

To generate biallelic *loxP* ESC clones, single guide RNA (gRNAs) targeting specific regions flanking *cis*-regulatory elements was designed using the CRISPR design tool (http://crispr.mit.edu/). gRNAs were cloned into the Cas9 expressing vector px459 v2.0 (Addgene #62988)[41,42]. Single strand DNA oligos were designed with ∼60 bp homology directed repair (HDR) arms flanking each side of the 34 bp *loxP* sequence and a restriction enzyme palindromic sequence (BamHI) for restriction digest genotyping of genomic PCR products. The *loxP* and restriction enzyme sequence was inserted between the PAM recognition sequence and the gRNA genomic targeting sequence. A single gRNA and single strand HDR oligo were co-transfected along with the gRNA (1-2 μg of each plasmid) into 1-2 × 10^6^ WT ESCs using Lipofectamine 2000 (Invitrogen #1168-019) in a single well of a 6 well plate. Transfected ESCs were selected with puromycin (2 μg/mL for two days only) and then passaged onto 10 cm at various dilutions and grown until single colonies appeared. Individual clones that were resistant to puromycin were isolated and expanded for genotyping. Primers designed outside of the HDR arms were used to genotype for enhancer deletion. Following genomic PCR, products were digested with XbaI (NEB, R0145s) for genotyping. Clones that demonstrate biallelic cutting were cloned into TOPO TA (Thermo Fisher #45-0641) for sequencing to confirm correct integration. *loxP* sequences (upstream or downstream of the targeted region) were inserted one at a time..

Cell lines described below were generated using the following CRISPR strategy. gRNAs were cloned into px459 v2. Plasmid was digested using Bbsl (NEB, R0539) and purified. gRNA oligos were phosphorylated and annealed using T4 PNK (NEB, M0201). The cut vector and annealed oligos were ligated overnight at 16°C. Ligated plasmids were transformed into NEB High Efficiency (NEB, C2987) bacteria, plated on LB+Amp plates and incubated overnight at 37°C. Colonies were picked and mini-prepped for sequencing, followed by maxi-preps once gRNA presence was verified. The same transfection protocol decribed above was followed for the following cell lines.

To generate a floxed −5 *Nanog* CRE ESC line for conditional deletion the 3’ *loxP* was inserted first using a gRNA and a HDR arm (Supplemental Table 1). The 5’ *loxP* sequence was subsequently inserted using using a gRNA and a HDR arm. The PAM sequences adjacent to the *loxP* sequences were mutated to prevent cutting of Cas9 following HDR. The gRNAs were used to constitutive delete the −5 CRE in [7]. ESCs were treated with 4OHT for 4 days at 2 uM to delete the −5 *Nanog* CRE.

To generate −5 *Nanog* CRE inverted clones we inserted the 5’ HDR oligo containing *loxP* in the opposite orientation into the ESC clone that contains the 3’ *loxP* above. The same 5’ gRNA above was used with a single strand HDR oligo. An ESC clone with *loxP* sequences in the opposite orientations was treated with 4-OHT at 2 uM for 3 days and cells were subcloned as described above. We confirmed biallelic inversion by genomic PCR with primers inside and outside of the *loxP* sequences. Clones that demonstrating wild-type, monoallelic, and biallelic orientation were cloned into TOPO TA plasmids, at least 4 individual clones were then isolated and sequenced to confirm correct integration.

To insert the −5 *Nanog* CRE downstream of the Nanog gene, a single gRNA was used to stimulate HDR of a modified version of pL451 (*loxP* sequence removed). The HDR vector contains a *Neomycin resistance* (*Neo*) cassette flanked *frt* sites and by homology regions (left arm chr6:122667133-122668329, 1197bp, mm9; right arm chr6:122668389-122669450, 1062bp, mm9). The left arm was cloned using KpnI and SalI and the right arm was cloned using BamHI and NotI. The enhancer (same sequence used in reporter assays in Blinka et al., 2016) was inserted adjacent to the left arm using SalI and EcoRI sites. The HDR plasmid was co-transfected along with a gRNA 5’ - TGGCTTGCATCCAATCTCTT - 3’ chr6: 122668369, mm9 (2-3 μg of gRNA and 6 ug of HDR plasmid) into 10 x 10^6 WT ESCs using Lipofectamine 2000. HDR vector arms and the enhancer were amplified off of a BAC [7] and fully sequenced in pBlueScript II SK(+) and matched the genomic reference sequence. Transfected ESCs were selected with puromycin (2 μg/mL first two days only) and G418 (350 μg/mL days 2-14) until single colonies appeared. Individual clones that were resistant to both puromycin and G418 were isolated and expanded for genotyping. *Neo* was removed by transfecting cells with a FLPe expressing plasmid driven by the CAG promoter. Primers designed outside of the HDR arms were used to genotype for enhancer insertion. Following genomic PCR to genotype, homozygous clones containing Neo were amplified and cloned into TOPO TA for sequencing to confirm correct integration.

Sub-enhancer deletions were generated using three gRNAs. Clones were genotyped using PCR primers designed around the sub-enhancers. Distance deletion clones were generated using four gRNAs. Clones were genotyped using PCR primers that surrounded the deleted portion.

All gRNAs and genotyping primers are listed in Supplemental Table 1.

### Generation of murine Nanogv5 Rescue Cell Line

The mouse Nanog sequence was synthesized by GeneArt Strings™ DNA Fragment (ThermoFisher). A C-terminus v5 tag was added to distinguish from endogenous Nanog protein. The synthesized DNA fragment was A-tailed and cloned into TOPO TA (Thermo Fisher #45-0641) to confirm the sequence. XhoI and NotI sites were designed at the 5’ and 3’ end of the DNA fragment so that it could be cloned into and the pPyCAG iH vector (hygromycin resistance, gifted from Austin Smith) for expression under a ubiquitous (CAG) promoter[13,40]. ESCs were electroporated with the linearized plasmid (Fsp1 NEB R0135) in the presence of hygromycin and individual clones were isolated as we have done previously and expanded for further experiments.

| Cell Lines in this paper |  |  |
| --- | --- | --- |
| Cell Line | Reference | Identifier |
| Mouse ESC floxed -5 SE with CreER<sup>T2</sup> at *Rosa26* locus | This paper | O6 |
| Mouse ESC floxed -5 SE with CreER<sup>T2</sup> at *Rosa26* locus and mNanogV5 at other locus | This paper | Clone 13 |
| Mouse ESC biallelically deleted -5 SE with CreER<sup>T2</sup> at *Rosa26* locus and mNanogV5 at other locus | This paper | Clone 13.5 |
| Mouse ESC -5 SE in inverted orientation with CreER<sup>T2</sup> at *Rosa26* locus | This paper | Q19.14 |
| Mouse ESC -5 SE closer to *Nanog* promoter, 2kb deletion between enhancer and promoter | This paper | 142 |
| Mouse ESC two copies of -5 SE inserted downstream of *Nanog* with CreER<sup>T2</sup> at *Rosa26* locus | This paper | R6.1 |
| Mouse ESC 5' sub-enhancer of -5 SE deleted with CreER<sup>T2</sup> at *Rosa26* locus | This paper | S20 |
| Mouse ESC 3' sub-enhancer of -5 SE deleted with CreER<sup>T2</sup> at *Rosa26* locus | This paper | T51 |
| Mouse ESC wildtype cells eGFP in Oct4 locus | Stelloh et al., 2016 | Oct4eGFPac |

### Total RNA RT-qPCR

Total RNA was harvested from cells following manufacturer’s protocol (TRIzol™ Reagent, Invitrogen #15596018). Genomic DNA was removed from total RNA samples using a DNA eliminator column step and passing RNA over a column following manufacturer’s protocol (RNeasy Plus Mini Kit, Qiagen#74134). Equal amounts of DNA-free total RNA were converted to cDNA using the iScript™ cDNA synthesis kit (Bio-Rad #1708891). Quantitative PCR (qPCR) was performed on a QuantStudio™ 6 Flex Real-Time PCR System (ThermoFisher). Quantifications were normalized to an internal control (Actin) for Reverse Transcriptase-qPCR (RT-qPCR) using the ΔΔCt method as we have done previously[7]. Primers used for RT-qPCR are described in Supplemental Table 1.

### Alkaline-Phosphatase staining

Bright field images and alkaline phosphatase staining were performed as previously described (Rao et al., 2010 and more, Sigma 86R-1kt). Briefly, cells were plated in 10cm dishes and treated for up to 6 days with vehicle or tamoxifen. Plates were rinsed 1x with PBS, fixed using citrate-acetone-formaldehyde fixative for 30s, rinsed with deionized water for 45s. Alkaline-dye mixture (diazonium salt solution + deionized water + Naphthol As-BI Alkaline Solution) was added to the plate and incubated for 15 minutes at room temperature in the dark. Dye mixture was removed from the plates, plates were rinsed for 2 minutes with deionized water and then air dried.

### Western Blots

Proteins were extracted in RIPA buffer and quantified as described in [43]. 10 µg of protein were loaded in each well of a gel (Bio-Rad # 567-1094, 567-1095, 456-1036). Blots were blocked in 5% milk/TBST for two hours at room temperature (RT). Primary antibodies to Nanog (Millipore; Cat # 5731) was used at 1:1000 or beta-Actin (Sigma; Cat # a5441) was used at 1:5000 in 5% milk/TBST for 90 minutes at RT. Blots were then washed and secondary antibody Donkey anti-Rabbit IgG-HRP (Santa Cruz; Cat # sc2313) was used at 1:5000 for 30 minutes at RT for Nanog. For beta-Actin, a secondary antibody (Santa Cruz; Cat # sc2064) Goat anti-Mouse IgM-HRP was used at 1:5000 for 30 minutes at RT. Antibody labeled proteins were detected using Amersham ECL Prime Western Blotting Detection Reagent (Cat # RPN2232).

### RNA-seq

RNA-Seq libraries were generated using the NEBNext Ultra™ RNA Library Prep Kit for Illumina (NEB #E7530). Libraries were quantified using the NEBNext Quant Kit (NEB #E7630), Agilent Tapestation 2200 (D1000 tapes) and run on a NextSeq 500, 36×36 PE. Library preparation and sequence were performed following manufacturer’s protocol. Data was analyzed using STAR (mm9)[44], Cufflinks[45] and DESeq[46] using default parameters through Basepair (www.basepairtech.com).

### Single-cell Analysis

Single cell analysis was performed using the Fluidigm C1 and BiomarkHD system following manufacturer’s protocol. Data was analyzed first using the Fluidigm Real-Time PCR Analysis Software to remove any data point with a poor melt curve or no amplification and then using R. Cts were normalized to *ACTB* measurements and any cell with an *ACTB* measurement above 8 was excluded as the quality of those samples could not be ensured. Data is represented as a difference from the Limit of Detection (as described by Fluidigm, SINGuLAR Analysis Toolset) and expressed as Log2Expression. Statistical difference was tested using a Mann-Whitney test with a p-value of 0.001.

### CUT&Tag

100,000 cells were collected and processed through the method described in Kaya-Okur et al., 2019, for Total RNA Polymerase II and RNA Polymerase II Ser5P (Cell Signaling Technologies, #54020). Libraries were quantified using the KAPA Quant Kit (#07960140001), Agilent Tapestation and sequenced on a NextSeq 500, 36×36 paired end. Data was processed as described in Kaya-Okur et al., 2019.

### ATAC-seq

ATAC-Seq libraries were generated as described previously on cells with the −5 SE floxed and cells with the −5 SE deleted, with Nanog expressed exogenously[39]. ESCs were plated 24h prior to experiment, collected and transposed for 30mins. Data was analyzed using bowtie2[47] using default parameters through Basepair (www.basepairtech.com).

### Data Set Reanalyses

All Chromatin Immunoprecipitation Sequencing (ChIP-Seq) and Global Run-on Sequencing (GRO-Seq) data sets displayed using the Integrated Genome Viewer (IGV) (data.broadinstitute.org). These data sets were previously downloaded and analyzed from the GEO[13]. Data sets are listed in Supplemental Table 2.

### Statistical analysis

Statistical analyses were done using Microsoft Excel and R. Statistical details of experiments can be found in the Fig legends. Two-tailed Student’s t-test comparisons (one sample and two sample t-test as indicated in the Fig legends) were performed and p-values < 0.05 were considered significantly different. Statistical significance was not shown for values within 20% of the control or between experimental values for RT-qPCR experiments as that is within the error of the assay. All error bars are standard error of the mean (SEM) between experimental replicates. For single-cell RT-qPCR, Mann-Whitney test was performed and p values < 0.001 was considered significantly different. Error bars are shown as standard deviations (SD). For primary transcript data, plasmid-based standards were used to quantify sample measurements. A RT-qPCR efficiency of >90% was set as a determinant of accuracy of an experiment. For all non-NGS methods a minimum of three replicates were done for all experiments.

Data is available on the GEO Omnibus (GSE143993) and can be accessed using reviewer token ehwfwcqqnlkhjah

## ACKNOWLEDGEMENTS

The authors would like to thank Drs. Emery Bresnick (University of Wisconsin); Erin M. Wissink and John T. Lis (Cornell University) for their assistance. This work was supported in part by funding from NIDDK (DK120152, DK10350) to PA and SB, respectively, and the MCW MSTP T32 (NIGMS, GM080202) to PA and SB. Additional support came from NIH (CA204231) to SR.

## AUTHOR CONTRIBUTIONS

PA, SB, and MR performed all experiments. KP and PA analyzed NGS data. PA, SB, MR and SR designed and interpreted experiments collectively. PA, SB, and SR wrote the manuscript with input from all authors.

## Supplemental Figure Legends

**Supplemental Fig 1.**
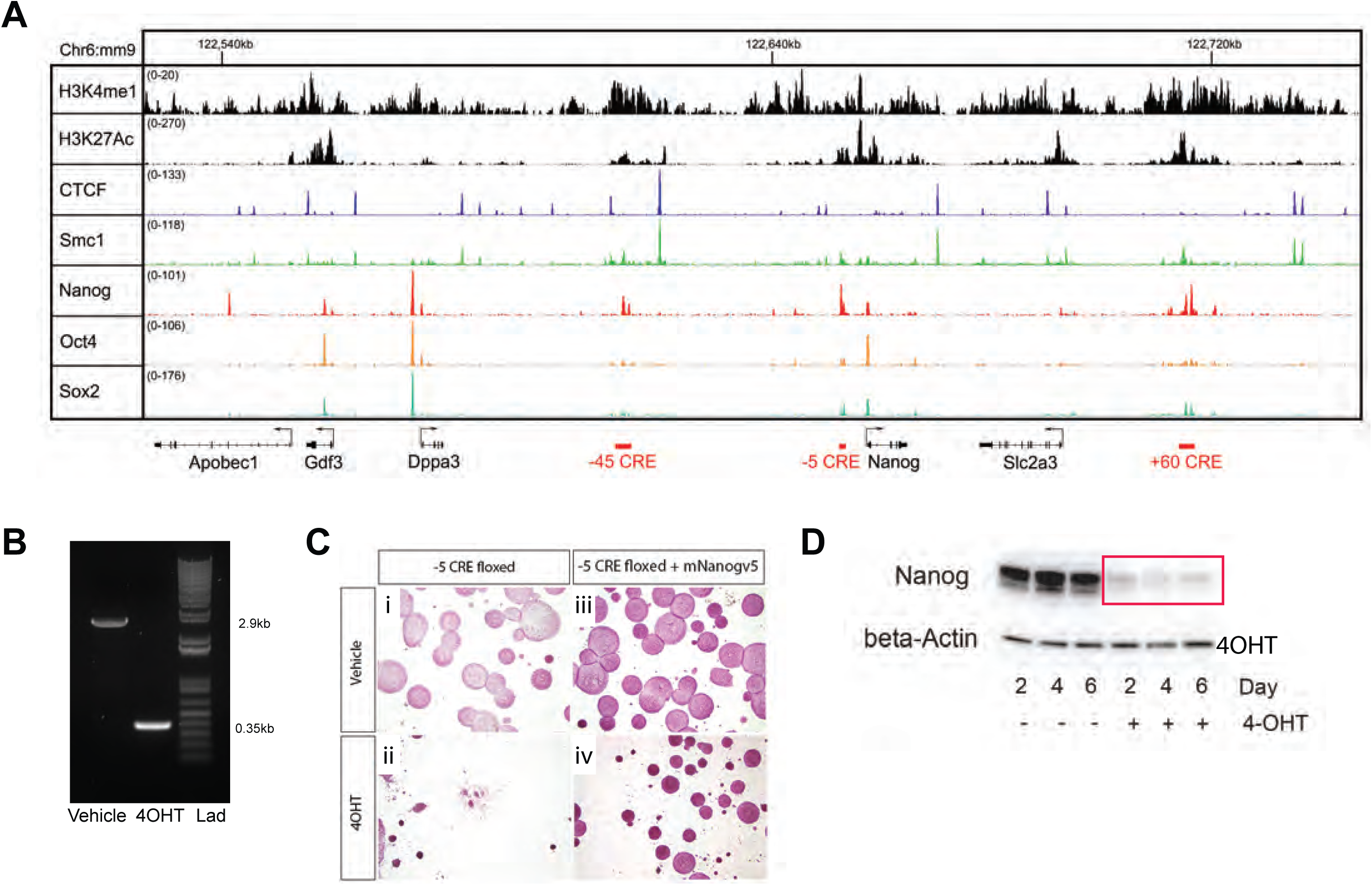
Nanog and the deletion of the −5 CRE. (A) IGV snapshot showing ChIP-seq tracks for H3K4me1, H3K27Ac, CTCF, Smc1, Nanog, Oct4 and Sox2. Genes and CREs are shown below the ChIP-seq tracks. X-axis is genomic position, Y-axis is normalized tag count. (B) PCR genotyping before and after tamoxifen (4OHT) treatment of cells depicted in Fig 1A and. (C) Alkaline Phosphatase of cells treated with tamoxifen (4OHT) – Fig 1A and E. (D) Western blot for Nanog in −5 CRE floxed cells at different days post 4OHT treatment. Actin is shown as a loading control.

**Supplemental Fig 2.**
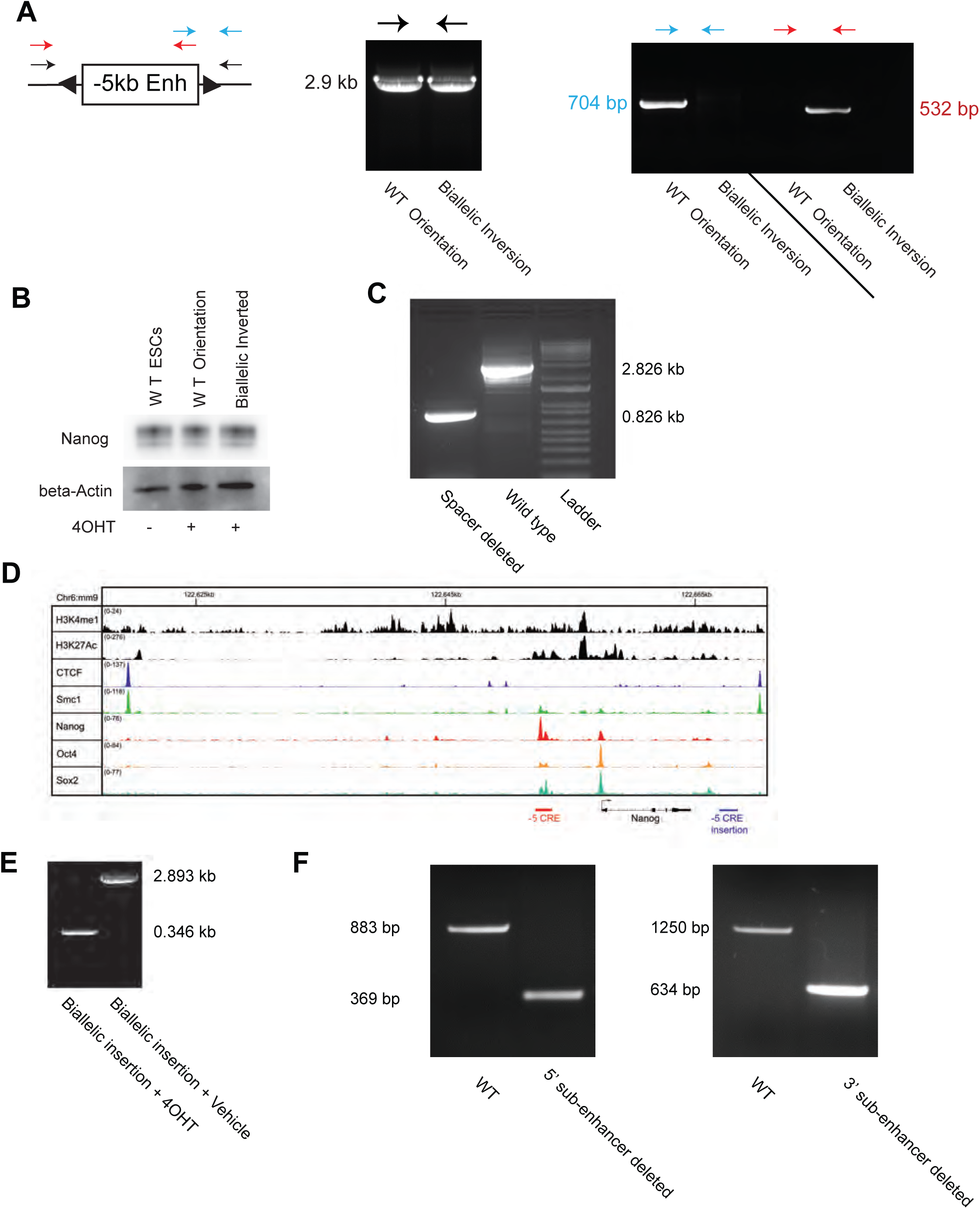
Manipulation of the −5 CRE. (A) PCR genotyping of −5 CRE inversion cells shown in Fig 3A. (B) Western blot of Nanog in −5 CRE cell lines with or without 4OHT treatment. Actin is shown as a loading control. (C) PCR genotyping of cell lines where the −5 CRE is closer to the *Nanog* promoter, as shown in Fig 3B. (D) IGV snapshot of −5 CRE downstream insertion. X-axis is genomic position, Y-axis is normalized tag count. (E) PCR genotyping for biallelic inserted cells before and after 4OHT treatment. F) PCR Genotyping of the two sub-enhancer deletions.

**Supplemental Fig 3.**
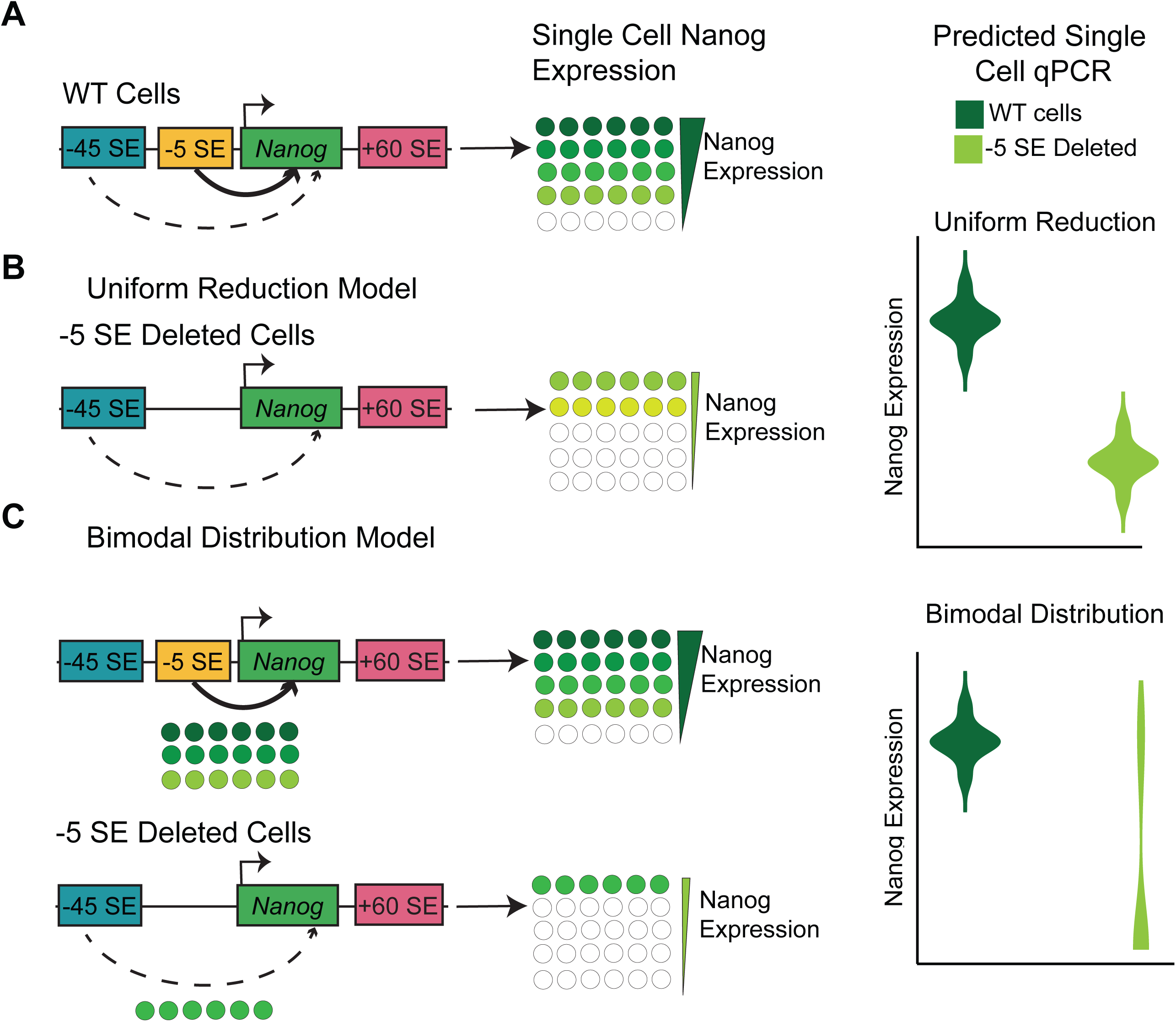
Predicted results of single-cell RT-qPCR based upon different models. (A) Wild-type cells and interaction between *Nanog* and the −45 SE and −5 SE, with the predicted distribution of *Nanog* expression across cells. (B) Model of changes in *Nanog* expression when there is a uniform reduction following enhancer deletion, implying the −5 SE is active in all cells. (C) In the bimodal distribution model, the −5 SE regulates more cells which have a higher level of *Nanog* expression cells, and upon deletion, the only cells that remain are ones that regulated by other enhancer that express lower levels of *Nanog* remain.

**Supplemental Fig 4.**
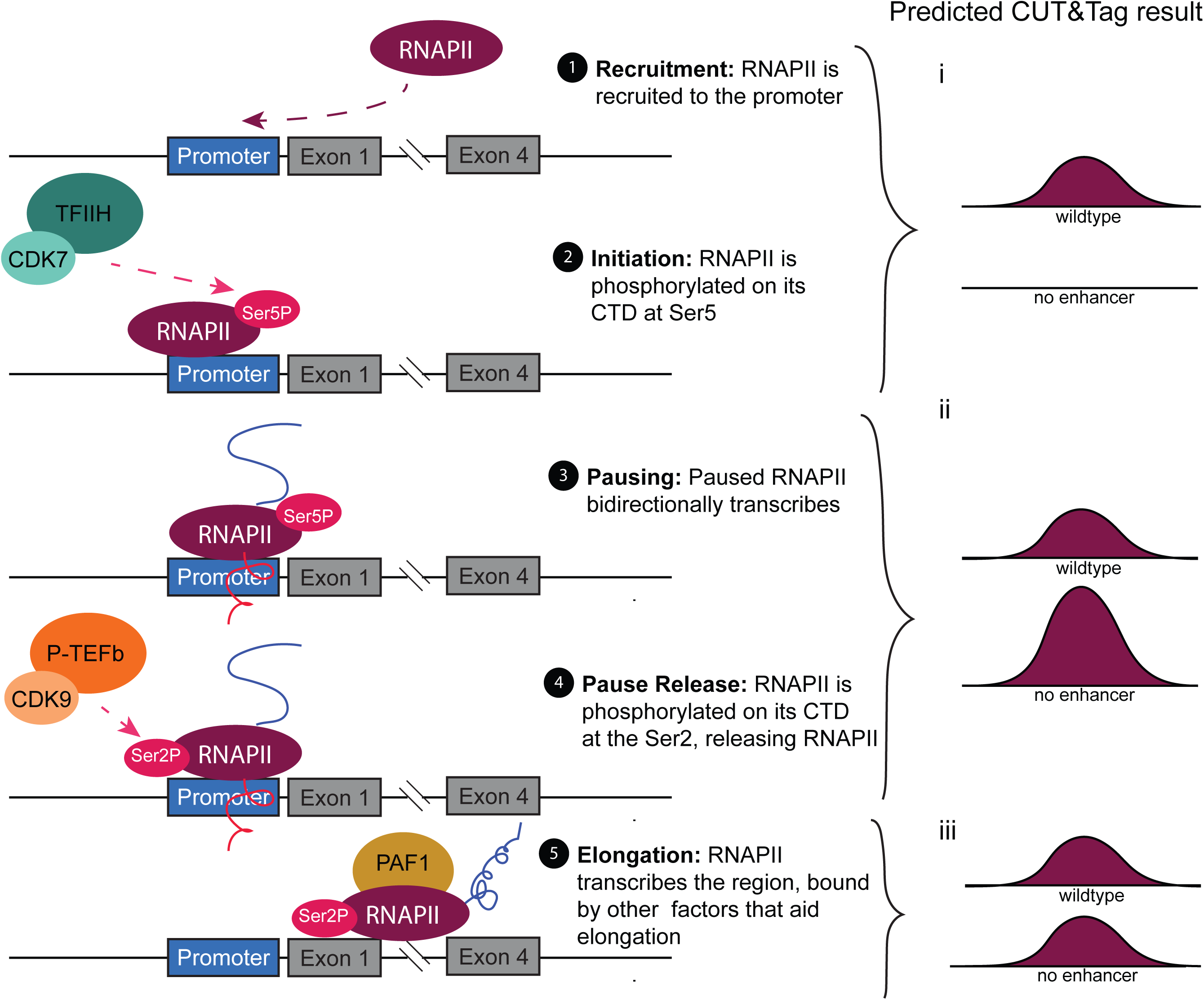
Enhancer-mediated gene expression. Overview of steps of transcription (1-5) and the predicted changes in Total RNAPII and RNAPII-Ser5P if an enhancer is regulating that step and is deleted (i-iii).

**Supplemental Table 1.**
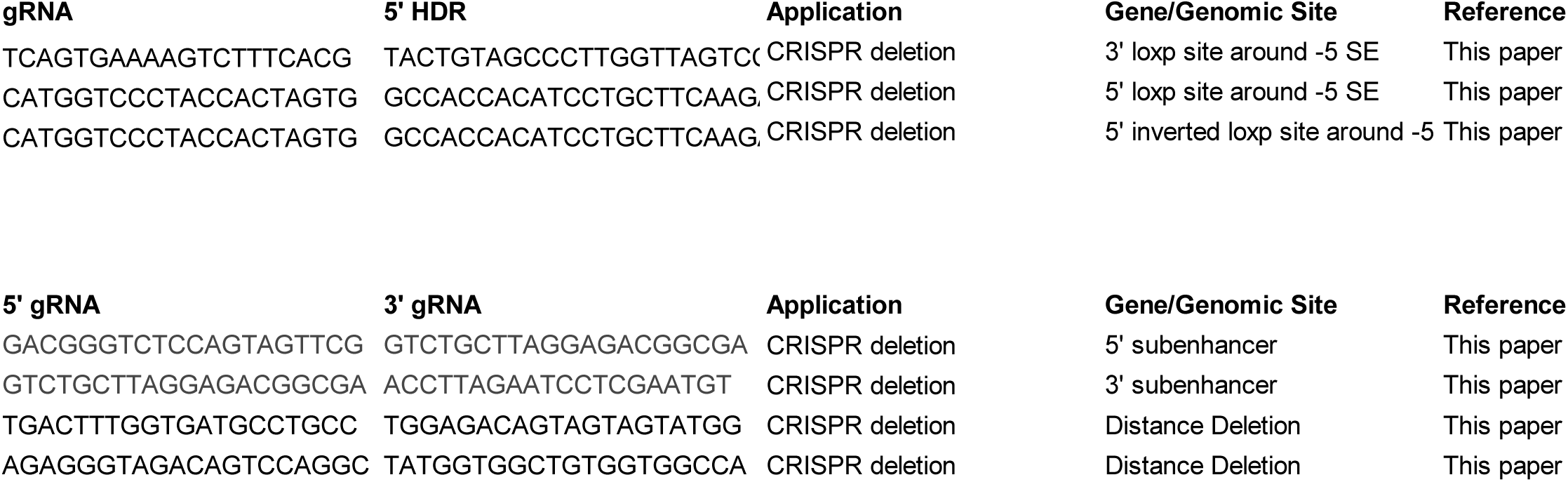
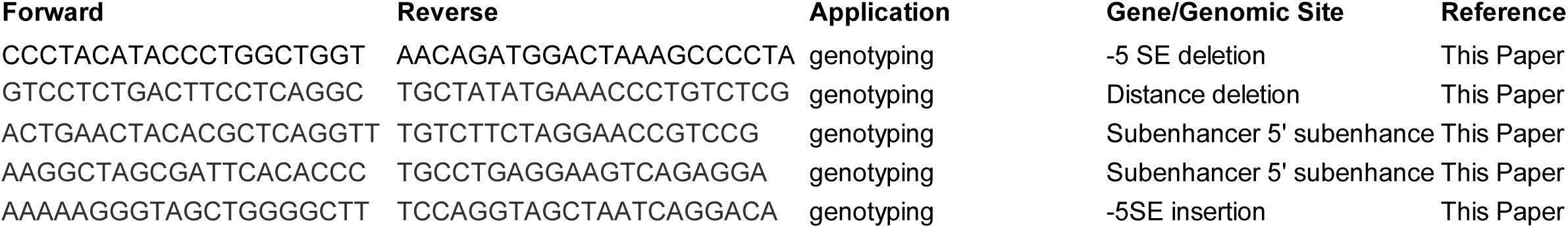
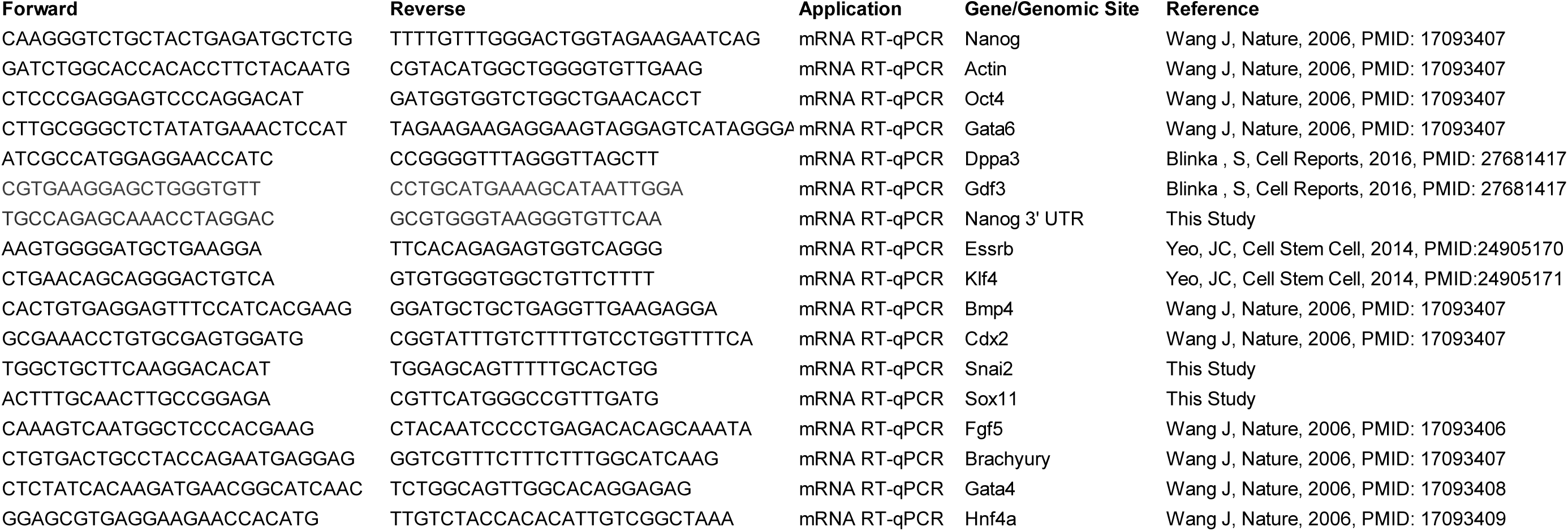
1) gRNAs for generating cell lines. 2) PCR genotyping primers for all cell lines generated in this paper. 3) primers for RT-qPCR

**Supplemental Table 2.** Accession numbers for published ChIP-Seq datasets analyzed.

**ChIP-seq datasets utilized in this study**
| **Protein/Histone** | **GSE** | **GSM(Experimental)** | **GSM (Control)** | **Reference** |
| --- | --- | --- | --- | --- |
| SMC1 | GSE22562 | GSM560341,GSM560342 | GSM560357 | Kagey MH *et al*, Nature, 2010. PMID: 20720539 |
| CTCF | GSE28247 | GSM699165 | GSM699168 | Handoko L *et al*, Nat Genet 2011. PMID: 21685913 |
| H3k4me1 | GSE24165 | GSM594577 | GSM594599 | Creyghton MP *et al*, PNAS. PMID: 21106759 |
| H3k27ac | GSE24165 | GSM594578,GSM594579 | GSM594599 | Creyghton MP *et al*, PNAS. PMID: 21106759 |
| Nanog | GSE11724 | GSM307140,GSM307141 | GSM307154,GSM307155 | Marson A *et al*, Cell 2008. PMID: 18692474 |
| Sox2 | GSE11725 | GSM307138,GSM307139 | GSM307154,GSM307155 | Marson A *et al*, Cell 2008. PMID: 18692474 |
| Oct4 | GSE11726 | GSM307137 | GSM307154,GSM307155 | Marson A *et al*, Cell 2008. PMID: 18692474 |

